# The *Aphelenchoides* genomes reveal substantial horizontal gene transfers in the last common ancestor of free-living and major plant parasitic nematodes

**DOI:** 10.1101/2022.09.13.507733

**Authors:** Cheng-Kuo Lai, Yi-Chien Lee, Huei-Mien Ke, Min R Lu, Wei-An Liu, Hsin-Han Lee, Yu-Ching Liu, Toyoshi Yoshiga, Taisei Kikuchi, Peichen J. Chen, Isheng Jason Tsai

**Affiliations:** Biodiversity Research Center, Academia Sinica, Taipei 11529, Taiwan; Genome and Systems Biology Degree Program, National Taiwan University and Academia Sinica, Taipei, Taiwan; Biodiversity Program, Taiwan International Graduate Program, Academia Sinica and National Taiwan Normal University, Taipei, Taiwan; Department of Life Science, National Taiwan Normal University, 116 Wenshan, Taipei, Taiwan; Department of Microbiology, Soochow University, Taipei, 111, Taiwan; Faculty of Agriculture, Saga University, Saga 840-8502 Japan; Department of Integrated Biosciences, Graduate School of Frontier Sciences, The University of Tokyo, Chiba, 277-8562, Japan; Department of Plant Pathology, National Chung Hsing University, Taichung, Taiwan

## Abstract

*Aphelenchoides besseyi* is a plant-parasitic nematode (PPN) in the Aphelenchoididae family capable of infecting more than 200 plant species. *A. besseyi* is also a species complex with strains exhibiting varying pathogenicity to plants. We present the genome and annotations of six *Aphelenchoides* species, four of which belonged to the *A. besseyi* species complex. Most *Aphelenchoides* genomes have a size of 44.7-47.4 Mb and are amongst the smallest in clade IV, with the exception of *A. fujianensis*, which has a size of 143.8 Mb and is the largest. Phylogenomic analysis successfully delimited the species complex into *A. oryzae* and *A. pseudobesseyi* and revealed a reduction of transposon elements in the last common ancestor of *Aphelenchoides*. Synteny analyses between reference genomes indicated that three chromosomes in *A. besseyi* were derived from fission and fusion events. A systematic identification of horizontal gene transfer (HGT) genes across 27 representative nematodes allowed us to identify two major episodes of acquisition corresponding to the last common ancestor of clade IV or major PPNs, respectively. These genes were mostly lost and differentially retained between clades or strains. Most HGT events were acquired from bacteria, followed by fungi, and also from plants; plant HGT was especially prevalent in *Bursaphelenchus mucronatus*. Our results comprehensively improve the understanding of horizontal gene transfer in nematodes.

## Introduction

The ability to parasitise plants has evolved in the phylum Nematoda on at least four occasions (Blaxter et al., 1998; Van Megen et al., 2009). The major plant parasites belonged to the Aphelenchodidae and Parasitaphelenchidae families making up the Aphelenchoidea superfamily and the Tylenchida order of clade IV nematodes (Bird, Jones, Opperman, Kikuchi, & Danchin, 2015); these plant parasitic nematodes (PPNs) collectively cause worldwide agriculture damages of over US$80 billion each year (Nicol et al., 2011). Root-knot nematodes in *Meloidogyne* genus cause the majority of these losses and were the first of PPNs to have their genomes sequenced (Abad et al., 2008; Opperman et al., 2008), followed by pinewood nematode *Bursaphelenchus xylophilus* (Dayi et al., 2020; Kikuchi, Cotton, Dalzell, Hasegawa, & Kanzaki, 2011), potato cyst nematode *Globodera pallida* (Cotton et al., 2014), soybean cyst nematode *Heterodera glycines* (Masonbrink et al., 2019) and others (Eves-van den Akker et al., 2016; Koutsovoulos et al., 2020; Wu et al., 2020). Comparing these genomes yield insight into several adaptions that allow PPNs to parasitize plants. Examples include effectors such as carbohydrate active enzymes (CAZyme), which are known to be secreted by PPNs and are hypothesized to be involved in degrading or modifying the composition of different plant structural tissues (Ali, Azeem, Li, & Bohlmann, 2017; Danchin, Guzeeva, Mantelin, Berepiki, & Jones, 2016). Some of these PPN-specific genes are known to be acquired from bacteria or fungi through horizontal gene transfer (HGT) (Jones, Furlanetto, & Kikuchi, 2005), giving nematodes the ability to adapt to different environments (Jones et al., 2005). Although numerous HGT genes have been identified and documented in different nematodes, research on the timing and subsequent maintenance of these genes, and why their copy numbers differ, has been restricted to a few PPN clades (Grynberg, Coiti Togawa, et al., 2020).

Currently, the only major groups containing plant parasitic nematodes that lack a reference genome are Trichodoridae and Aphelenchoididae. Of particular interest is *Aphelenchoides besseyi*, a foliar nematode that infects almost 200 plant species in 35 genera (Jen, Tsay, & Chen, 2012). This nematode is 0.4-0.8 mm in body size, has a life cycle around 10 to 12 days and it can reproduce in extreme environments, making it hard to eliminate. Better known as the rice white tip, *A. besseyi* infects important agronomic crops such as rice, soybeans and strawberries (Oliveira et al., 2019; Subbotin et al., 2020), causing necrosis and distortion of its host’s leaves (Jen et al., 2012; Oliveira et al., 2019; Wang et al., 2014). The nematode has reportedly been responsible for up to a 60% crop loss in some cases (Koenning et al., 1999; Lilley, Kyndt, & Gheysen, 2011) and was listed among the top ten plant parasitic nematodes in a 2013 review (Jones et al., 2013). Despite the economic damage these parasitic nematodes inflict, particularly in Asia, little is known about the basic biology, genetic diversity or evolution of *A. besseyi* and other Aphelenchoididae members.

The taxonomic status of *A. besseyi* was reevaluated several times before molecular data were available. After its first description by Christie (1942), Yokoo (1949) identified another *Aphelenchoides oryzae* species, but this was later considered a synonym of *A. besseyi* due to the overlaps of several morphological characters (Allen, 1952). Several characteristics such as stylet length (Fortuner, 1970) have been added as standards to identify *A. besseyi* globally (Bulletin, 2017). More recently, molecular analyses indicated that *A. besseyi* may consist of several genetically diverged clades.

We previously identified copy number variations of genes encoding cell-wall-degrading enzymes including glycosyl hydrolase family 5 (GH5) and GH45 cellulases between *A. besseyi* of different host origins (Wu, Kuo, Tsay, Tsai, & Chen, 2016). An 18S phylogeny separated these strains isolated from rice and fern unambiguously, and more recently new literature identified variations in different molecular markers in different hosts that are original to *A. besseyi* (Oliveira et al., 2019; Xu et al., 2020). Subbotin *et al*. used a combination of molecular makers (28S, ITS and mitochondria *COI* gene) (Subbotin et al., 2020) to reclassify foliar nematodes into three distinct clades: *A. besseyi* isolated mainly from strawberries, *A. oryzae* isolated mainly from rice and *A. pseudobesseyi* from wood fern. This study proposed this *A. besseyi* is a species complex, with each cryptic species associated with, but not restricted to, particular plant hosts. It has been reported that *A. besseyi* isolated from different hosts have varying levels of pathogenicity. For instance, *A. besseyi* populations isolated from strawberries were unable to parasitise rice (Koenning et al., 1999), while populations of this species from bird’s-nest fern were able to reproduce in both rice and strawberries (Wang et al., 2014). From an evolutionary perspective, *A. besseyi* is also interesting because its primitive plant parasitism was a relatively recent evolutionary adaptation (Rybarczyk-Mydłowska et al., 2012).

In this study, we sequenced and annotated the genomes of four *A. besseyi* strains isolated from different plants which we later designated as *A. pseudobesseyi* and *A. oryzae*, and another two species in the Aphelenchoididae family (*Aphelenchoides bicaudatus* and *Aphelenchoides fujianensis*). We compared the proteomes of six Aphelenchoididae members with 21 other representative nematodes to delimit species relationships and investigated their gene family dynamics. We identified synteny with representative nematodes and inferred rearrangement events to determine how the three chromosomes of *A. besseyi* was evolved. The availability of the *Aphelenchoides* assemblies allowed us to systematically determine the horizontal gene transfer-acquired genes in nematode genomes. By inferring the evolutionary origins of these HGT genes we found historical HGT events that shaped nematode evolution.

The major event occurred in the last common ancestor of clade IV nematodes and may have contributed to the early adaptation of these nematodes.

## Results

### Genome assemblies and annotations of six *Aphelenchoides* species

We sequenced and assembled the genomes of six nematodes in the *Aphelenchoides* genus (four *A. besseyi*, one *A. bicaudatus* and one *A. fujianensis*). These species were chosen to represent the Aphelenchoididae family and *A. besseyi* strains isolated from three plant hosts (**table S1**) to delimit their relationship within the species complex. For each species, an initial assembly was produced from either 70-148X Oxford Nanopore or 113-422X Pacbio reads using Flye assembler (Kolmogorov, Yuan, Lin, & Pevzner, 2019) and further polished using Illumina reads (**table S2**).

Among *A. besseyi* assemblies, the VT strain isolated from tape grass *Vallisneria spiralis* had the highest genome quality with N50 5.4 Mb (hereafter denoted as APVT). The contigs of this strain were further scaffolded with 150X Hi-C reads using the Juicer program (Durand et al., 2016) (**fig. S1**) yielding a final assembly of 44.7 Mb (N50 = 16.9 Mb). More than 99% of this assembly was in three scaffolds, presumably corresponding to three chromosomes (Yoshida, Hasegawa, Mochiji, & Miwa, 2009) (2n=6). Five *Aphelenchoides* assemblies ranged from 44.7 to 47.4 Mb (N50 = 12.2-17.8 Mb; **table S3**), and a sixth assembly (*A. fujianensis*) of 143.8 Mb (N50 = 553 kb; **table S3**) was estimated to be triploid (Ranallo-Benavidez, Jaron, & Schatz, 2020) (**fig. S2**). Although not present in the assemblies, the telomere motif TTAGGC was identified in the reads of *A. pseudobesseyi* at low coverage (**Supplementary Info**), which is consistent with the sister group species of *B. xylophilus* (**table S4**) and indicates the presence of telomeres in these species.

Using the proteomes of *Bursaphelenchus xylophilus* and *Caenorhabditis elegans*, and the transcriptomes of pooled worms in each species as evidences, we predicted 11,701 to 12,948 protein-coding genes in six *Aphelenchoides* species with Maker2 (Cantarel et al., 2008) (**table S3**). With the exception of *A. fujianensis*, these were fewer protein coding genes in these species than in Tylenchida nematodes (12,762 to 19,212) and free-living nematodes (20,184 to 20,992). The completeness of annotated genes was estimated to be 76.4–81.3% based on a BUSCO assessment, lower than that of *Bursaphelenchus* species (83.0–89.4%), but higher than that of Tylenchida (59.8–75.4%) nematodes. 66.5% to 71.0% of genes in *Aphelenchoides* could be assigned at least a domain from the Protein family (Pfam) database (Finn et al., 2014). In addition, orthologous groups were inferred with the proteomes of six *Aphelenchoides* with 21 other nematodes using Orthofinder (Emms & Kelly, 2019). With the exception of *A. fujianensis*, 78.5–85.4% (*A. fujianensis* = 48.7%), 69.4–76.9% (*A. fujianensis* = 42.8%) and 87.5–98.7% of *Aphelenchoides* genes were orthologous to *B. xylophilus, C. elegans* and at least one other nematode species, respectively, suggesting that the reduced proteome in most *Aphelenchoides* was mainly comprised of conserved genes among nematodes.

### Phylogenomics delimit species complex of *Aphelenchoides besseyi*

To investigate the evolution of plant-parasitic nematodes and the relationships among members in the *A. besseyi* species complex, a maximum-likelihood phylogenetic tree was constructed based on 75 low-copy orthologues. The phylogeny is consistent with the previous study (Kikuchi, Eves-Van Den Akker, & Jones, 2017): the major plant parasitic nematodes were divided into Aphelenchoidea and Tylenchida, and six *Aphelenchoides* species were grouped as sister to *Bursaphelenchus* (**fig. 1a**). The *A. besseyi* strains were clustered into two groups based on their hosts, suggesting that relationships in these species within the *A. besseyi* species complex were associated with host preferences. Combined with the previous 28S phylogeny of the *A. besseyi* species complex (Subbotin et al., 2020) (**fig. S3**), we further designated these two groups as *A. oryzae* and *A. pseudobesseyi* groups isolated from rice or other plants (land grass and bird’s-nest fern). The median nucleotide and amino acid identity was 86.6% and 90% between these two groups, respectively (**fig.S4**). Such levels of divergence were comparable to species comparisons of other clades (87.2% and 92.8% amino acid identity in *B. xylophilus–B. mucronatus* and *M. incognita–M. floridensis*, respectively). Strains in each group also differed in heterozygosity (0.017-0.019% in *A. oryzae* vs 0.071-0.075% in *A. pseudobesseyi*) and changes in recent effective population sizes inferred using pairwise sequentially Markovian coalescent (PSMC) analysis (Nadachowska-Brzyska, Burri, Smeds, & Ellegren, 2016) (**fig.5**). Together these results demonstrated that relationships among the two designated clades in the *A. besseyi* species complex were distinct species, well-differentiated at the genome level, despite being challenging to differentiate based solely on morphology (Subbotin et al., 2020).

**Figure 1.**
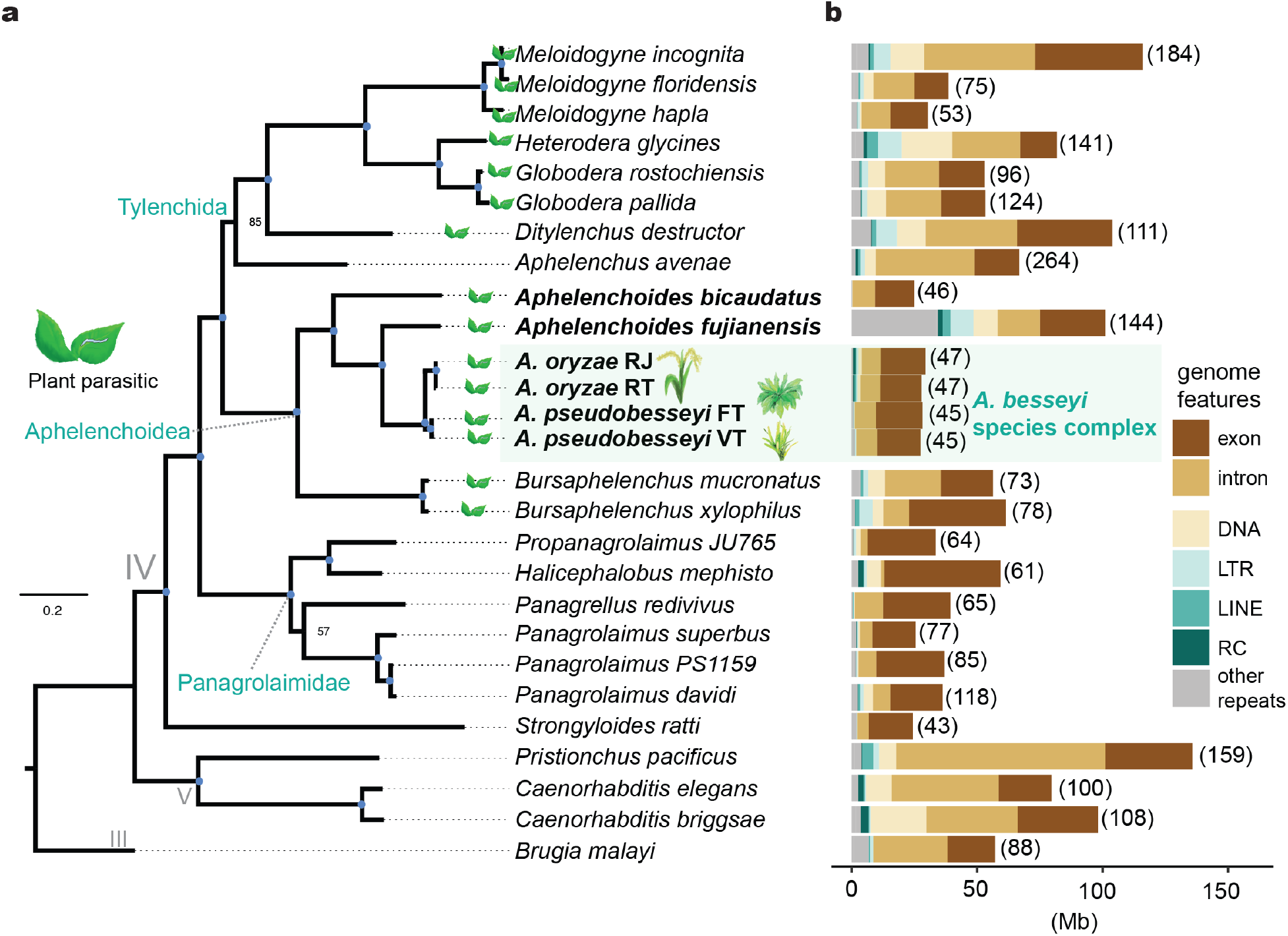
Phylogenomic analysis of plant-parasitic nematodes. **a**. The phylogeny of 27 representative nematodes was inferred based on the concatenation of protein sequences from 75 low-copy orthologues. The blue dots in branches denote a bootstrap value of 100. **b**. The size of the genome features in nematodes, repeats containing DNA transposons (DNA), long interspersed nuclear elements (LINE), long terminal repeats (LTR), rolling-circular (RC) and the unclassified repeats are labelled as “other repeats”. Numbers in brackets denote genome assembly sizes in megabases.

### Most *Aphelenchoides* species have reduced genomes and repetitive elements

The *Aphelenchoides* genomes were smaller than those of other plant-parasitic nematodes (**fig. 1b** and **table S3**). Much of the reduction can be explained by the reduced markup of repeat content compared to other nematodes (**fig. 1b**). The dominant transposable elements of *Aphelenchoides* were DNA transposons—which were reduced in content (0.14–1.36 Mb vs. 4.2–22.1 Mb in other nematodes)—and number of families (1–7 in *Aphelenchoides* compared to 9 and 26 in *B. xylophilus* and *H. glycines*, respectively) compared to other nematodes. Fewer LTR (0.07–0.8 Mb vs. 0.24–9.3 Mb) and LINE (0.0006–0.66 Mb vs 0.02–4.5 Mb) retrotransposons were also observed in this genus. These results suggest that the reduced genome sizes in *Aphelenchoides* might have been caused by the loss of transposable elements and led to the eventual loss of entire families in some cases (**fig. 2a**). Within the *A. besseyi* species complex, *A. pseudobesseyi* contained significantly fewer DNA transposons, LTR and LINE retrotransposons than *A. oryzae* (**fig. 2a and fig.6**).

**Figure 2.**
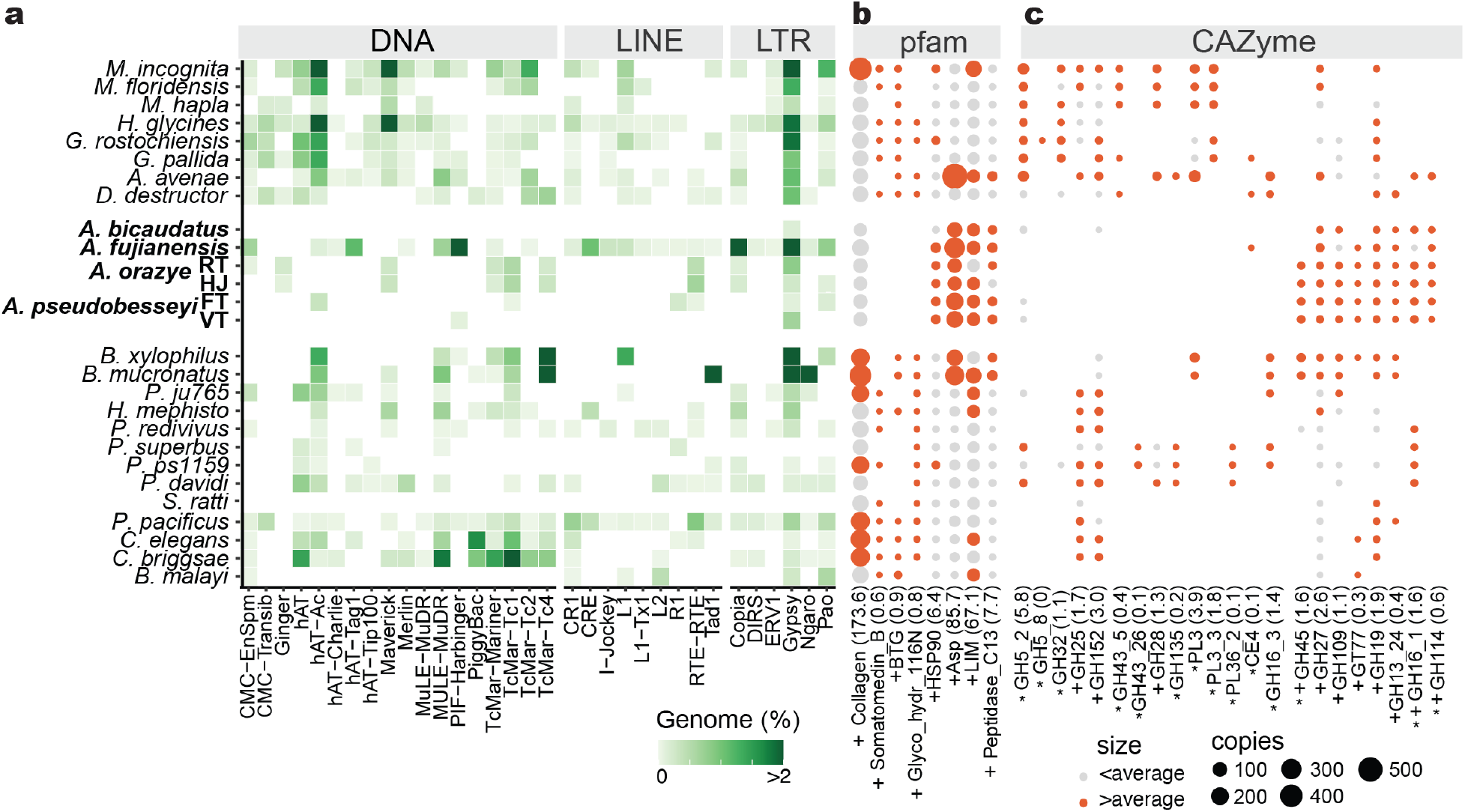
Repeat and proteome contents in nematodes. **a**. The genome proportions of DNA, LINE and LTR transposable elements in nematodes shown by genome percentage **b-c**. Protein families and CAZyme gene copy numbers vary significantly among nematodes. The dot size represents the copy number of each domain and the different colour represents the copy number of domains larger or lower than average copies shown in brackets. Pfam and CAZyme families that were significantlyy different between *Aphelenchoides* species and other nematodes are denoted by the “+” symbol. CAZyme families that was identified as having been acquired horizontally are denoted by the “*” symbol.

### Gene family specialization in the *Aphelenchoides* species

To gain insight into the genomic features associated with the biology and plant parasitism of *Aphelenchoides*, we aimed to identify expansion or contraction of protein domains and carbohydrate-active enzymes (CAZymes) specific to this clade. We observed 66 enriched and 31 reduced protein domains in the four members of the *A. besseyi* species complex compared to 21 other nematodes. (Wilcoxon rank-sum test q value < 0.05; **fig. 2b** and **table S5**). Reduced domains included collagen (90–109 copies in the *A. besseyi* species complex vs. 72–407 in others), Somatomedin B and BTG/Tob (Winkler, 2010). Genes containing collagen domains were reportedly associated with capsule formation; the reduced copy of collagen domains in *Trichinella spiralis* were thought to contribute its lower host-specificity than other nematodes (Mitreva et al., 2011), and may be related to the wide host range of *A. besseyi*. In contrast, Aphelenchoidea members possess on average four-fold (91–314 vs. 4–555 copies) more aspartic proteases (ASP) than other nematodes (**table S5**). Other expansions included LIM and peptidase C13 domains, which participate in participating in the regulation of cell motility and cell growth (Koch, Ryan, & Baxevanis, 2021) or degradation of protein tissues in a host (Dall & Brandstetter, 2016), emphasizing that these domain dynamics are associated with adaptations to plant parasitism.

The plant cell wall acts as a primary defensive barrier and the production of CAZyme families are important for PPNs to infect plants. A total of 132 CAZyme families were identified in the representative 27 nematode species. Of these, 59–67% of the CAZyme families were observed in Aphelenchoidea which is similar to the 55–66% and 58-68% of the families in Tylenchida and free-living nematodes (**table S6**), respectively. A total of 13 families were significantly expanded or lost in the *Aphelenchoides* genus **(fig. 2c**), including GH16, GH27 and GH45. GH16 serves as the putative β-glycanases involved in the degradation or remodelling of cell wall polysaccharides (Holm Viborg et al., 2019), GH16 had one to six copies in Aphelenchoididae nematodes and was not identified outside this clade except in *D. destructor*, in which there were three copies. There are three to 11 copies of GH27— which are reportedly involved in the function of hemicellulose and associated with α-galactosidase activity in both bacteria and fungi (Coconi Linares, Dilokpimol, Stålbrand, Mäkelä, & de Vries, 2020)— in Aphelenchoidea, but fewer in the Tylenchida nematodes. The previously identified GH45 present in Aphelenchoidea nematodes (Wu et al., 2016)—involved in the degradation of beta-1,4-glucans in the plant cell wall (Wang et al., 2014)—possess different copy numbers between *A. pseudobesseyi* and *A. oryzae* and were absent in *A. fujianensis* and *A. bicaudatus*, suggesting differential maintenance of these genes in the same genus may have contributed to variations of pathogenicity to plants.

### Chromosomal evolution of PPNs

To investigate the extent of the karyotype rearrangements in *Aphelenchoides*, we inferred the synteny relationships among *A. pseudobesseyi* (chromosome n=3) (Yoshida et al., 2009), *B. xylophilus* (n=6) and *C. elegans* (n=6) using single copy orthologs. Within the three *A. pseudobesseyi* chromosomes, orthologs belonging to all *C. elegans* chromosomes were clustered into distinctive blocks (**fig. 3a**) suggesting a fusion of ancestral chromosomes. These regions remained contiguous and contained 148-801 orthologous genes that could be assigned from individual chromosomes presumably not yet broken down yet by recombination, allowing us to pinpoint the fusion points and infer the order of rearrangement events based the constitution of chromosomes (**fig. 3b**). We encountered instances of where an ancestral chromosome was found in different parts of the *A. pseudobesseyi* chromosomes, suggesting fission also took place. In the case of chromosome IV—which remained homologous in *C. elegans* and *B. xylophilus*—corresponding synteny blocks in *A. pseudobesseyi* were identified in the arm of chromosome 2 and chromosome 1 separated by regions of chromosome III origin (**fig. 3b**). The majority of the ancestral sex chromosomes were unambiguously assigned to chromosome 2, and remapping of male sequences showed equal coverage along the chromosomes (**fig.S7**), suggesting that the Aphelenchoidea superfamily including *A. besseyi* exhibited a stochastic sex determination system that was recently characterized in *B. xylophilus* (Shinya et al., 2022). Within the *A. besseyi* species complex, a total of 91% and 88% of genomes were in synteny between APVT and AORT, respectively. Intra-chromosomal inversions were common at chromosome arms. In addition, we identified a major inversion of length 3.4 Mb long located in the centre of chr 2 (**fig. 3c**) suggesting rearrangement is still ongoing. Both the LTR and LINE retrotransposons were enriched in the chromosome arms of the *A. oryzae* strain (AORT) (**fig. 3c and fig.S6**), which is consistent with repetitive element distributions in other nematode chromosomes (Woodruff, 2019). In contrast, only the LTR retrotransposons were found in the two chromosome arms of *A. pseudobesseyi*, suggesting that these repeats were differentially maintained after speciation.

**Figure 3.**
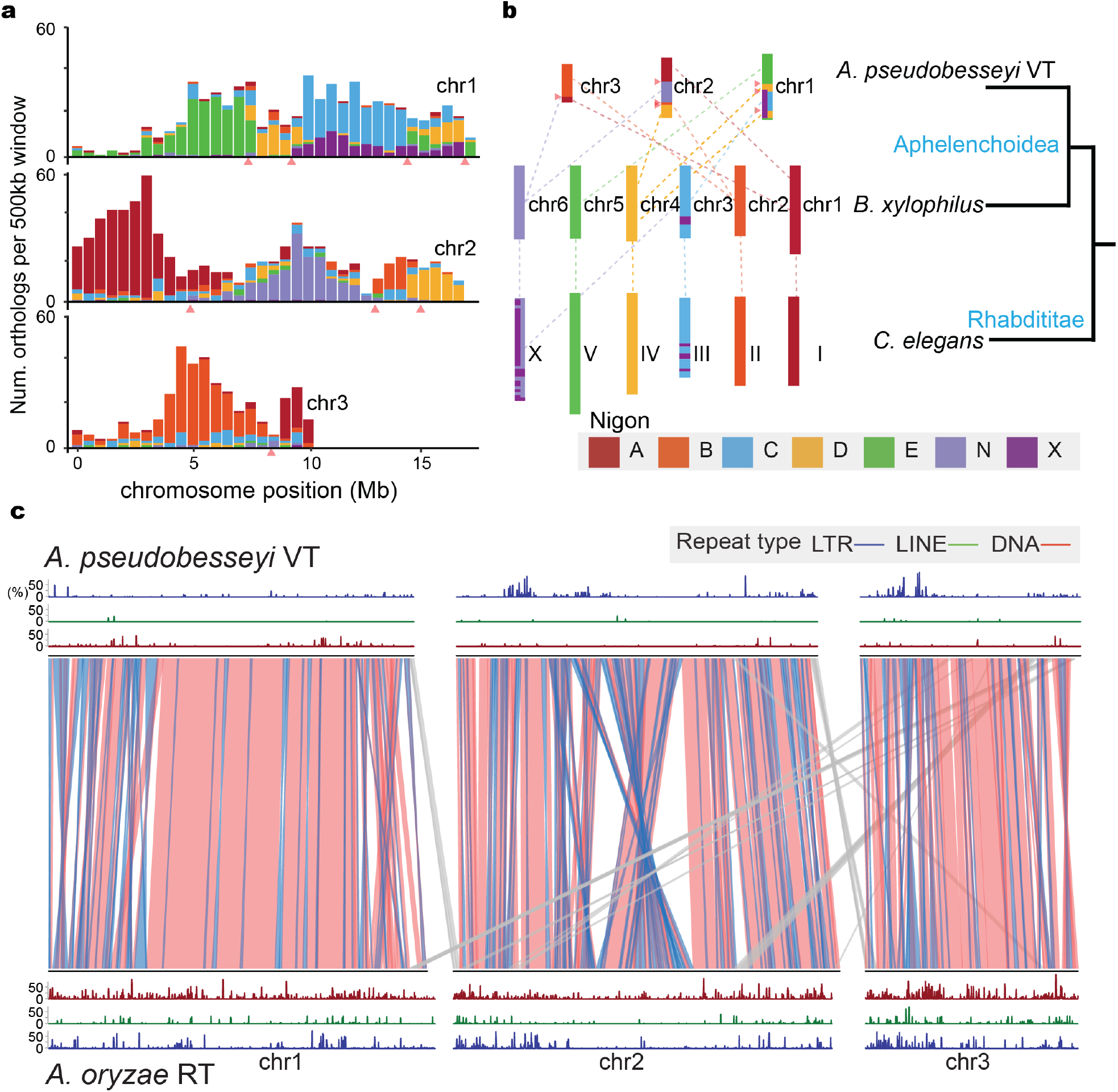
Chromosome evolution of plant-parasitic nematodes. **a**. The density of pairwise single-copy orthologs between *A. pseudobesseyi* VT and *B. xylophilus*. Colours denote the Nigon elements inferred from 15 rhabditid nematodes (de la Rosa et al., 2021) and the putative chromosome fusion sites in APVT are labelled with red triangles. **b**. Dashed lines indicate rearrangement events. **c**. The synteny relationship and the distribution of transposable between *A. pseudobesseyi* VT and *A. oryzae* RT. Blocks indicate synteny links between the two strains, and the line colours correspond to inversion (blue) and inter-chromosomal rearrangement (grey). Distribution of DNA transposons (red), long interspersed elements (green) and long terminal repeats (blue) between two stains are shown.

### Major episode of HGT in clade IV nematodes

In plant parasitic nematodes, the GH5 cellulase was found present in Tylenchida and only *A. pseudobesseyi* and *A. bicaudatus* within the Aphelenchoidea clade (Danchin et al., 2010; Wu et al., 2016), raising the possibility that many of the horizontal gene transferred (HGT) genes were acquired in the last common ancestor of major PPNs but were differentially lost. To identify such events, a total of 27 proteomes from representative nematodes including the *Aphelenchoides* genomes were searched for evidence of HGT by calculating the Alien Index (AI) score using Alienness (Rancurel, Legrand, & Danchin, 2017), which has been applied to infer HGT candidates in various nematode species (Grynberg, Togawa, et al., 2020; Schiffer et al., 2019; Siddique et al., 2021). We identified a total of 1,675 HGT orthogroups in 22 nematodes. Placing these orthologs designated as events onto the species phylogeny assuming a parsimonious scenario (Campoy & González-Martín, 2017), indicated that HGT started in the last common ancestor of clade IV nematodes (**fig 4a**). Examples include GH16, GH32, GH43 and the aforementioned GH5 cellulases. We inferred a total of 161 orthogroups were acquired in this episode, and most of their origins were inferred to be bacteria (78.3%) (**table S7**) belonging to different genera, suggesting multiple acquisitions took place. Of these, we found 36 Pfam terms such as ABC transporter that were identified in multiple orthogroups suggesting some convergence in the acquired functions (**table S8**).

**Figure 4.**
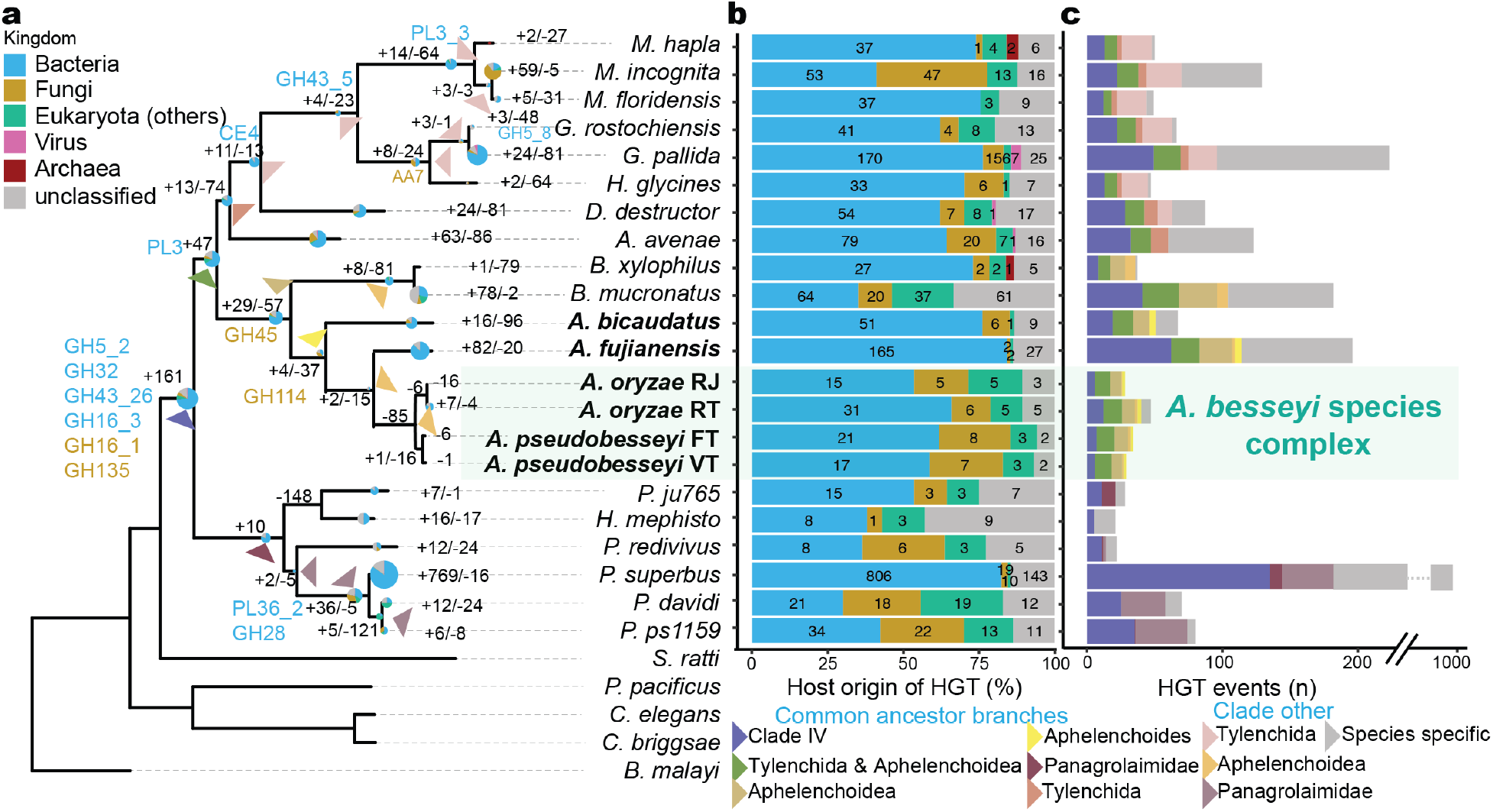
The evolution of genes acquired from horizontal gene transfer (HGT) in clade IV nematodes. **a**. HGT orthogroups were inferred by the AI score (Rancurel, Legrand, & Danchin, 2017) > 0 across 27 representative nematodes; the HGT families gained or lost are shown in the branches. Horizontally acquired CAZymes are annotated. The proportions of donor origins in each HGT orthogroup belonging to different kingdoms of donors are shown as pie charts. The size of the pie chart corresponds to the total number of HGT orthologues in branches; the chart was normalized using: log_5_(total number). **b**. The distribution of HGT families transferred from different kingdoms, with the same denoted color scheme as same as figure a. **c**. Number of HGT events among different inferred time points corresponding with the branches which are marked with triangles in the phylogeny.

The revised GH5 cellulase phylogeny indicated an ancient duplication took place before the divergence of PPNs (**fig 5a**). One clade contains orthologs of the three *Panagrolaimid* (*P*. sp. PS1159, *P. superbus* and *P. davidi*), and Tylenchida, and the other clade contains members of Aphelenchoidea and Tylenchida nematodes, which emphasises that the fate of the HGT genes was governed by duplications and loss.

**Figure 5.**
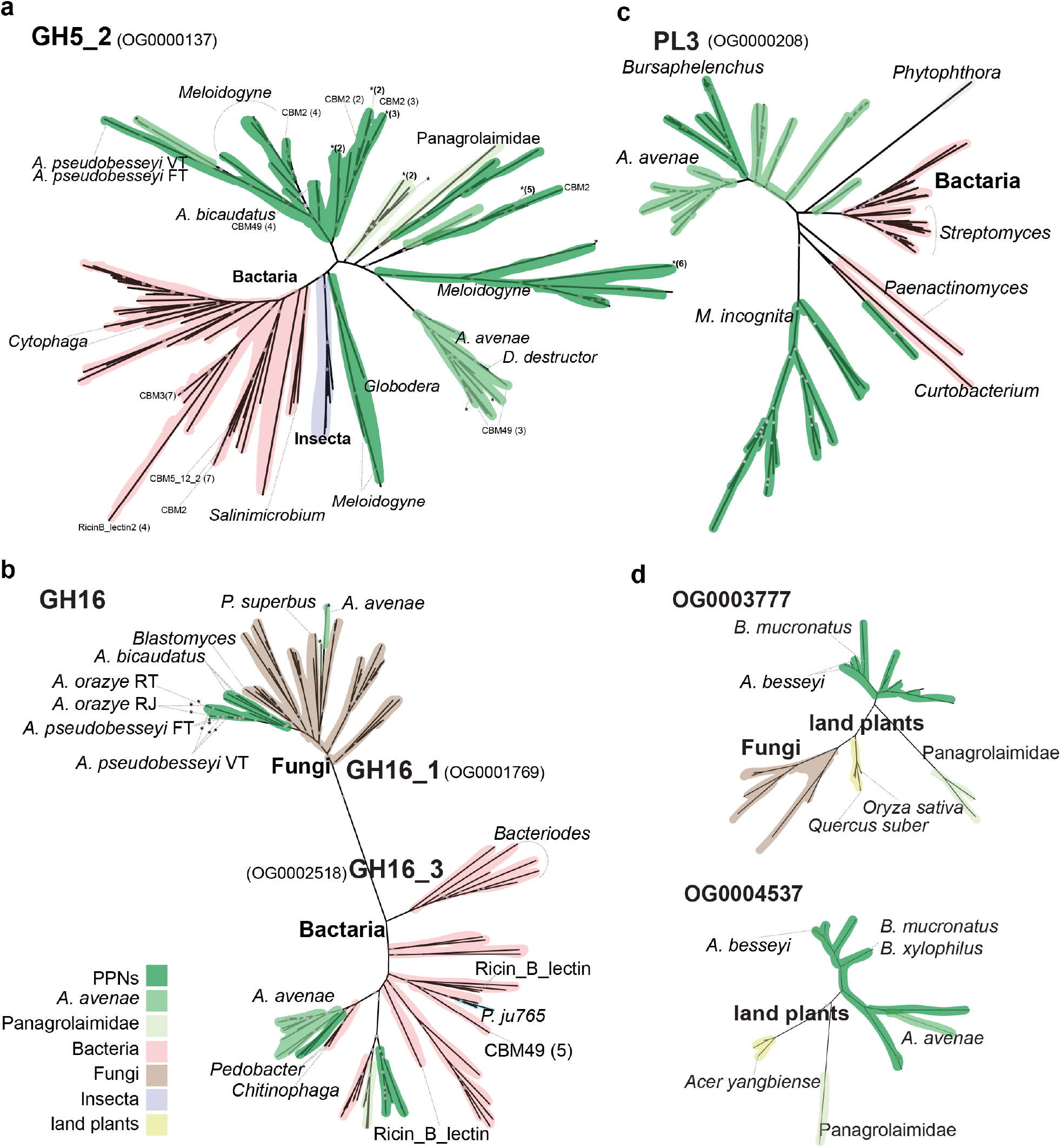
Phylogenies of nematode HGT orthogroups and their donor origin. **a**. GH5_2 **b**. GH16 **c**. PL3 **d**. OG0003777 and OG0004537. Different colours denote different kingdoms and species as shown in legend. Nematode gene copies with negative AI values were marked with an asterisk. Additional Pfam domains are labelled when available. Nodes with iqtree UFBoot and SH-aLRT bootstrap support > 80% are labelled as grey circles.

Interestingly, the closest GH5 bacterial orthologs were *Salinimicrobium xinjiangense* and *Leeuwenhoekiella sp*., which belonged to Flavobacteriaceae family and were from marine environments. We observed two GH16 subfamilies in nematodes. GH16_3 in Tylenchida and *Bursaphelenchida* nematodes were clustered with bacterial origin sequences, whereas GH16_1 of *Aphelenchoides* and *Panagrolaimus* nematodes were clustered with fungal origin (**fig 5b**), suggesting that the two GH16 groups arose independently. GH32 in *G. pallida* (Danchin et al., 2016) is believed to play a role in the function of fructose hydrolysis and was found in one *Panagrolaimus* in addition to several Tylenchida nematodes (**fig.S8**). GH43 was identified at two distinct clusters of bacterial origin in Tylenchida and *Panagrolaimid* nematodes which have been proposed to be involved in degradation of the hemicellulose in plants (Morais et al., 2021) (**fig.S9**).

The next major episode of acquisition took place in the common ancestor of PPNs, with 47 orthogroups (**fig. 4a)**. These families included pectate lyases 3 (PL3) which is associated with cell wall degradation (Atanasova et al., 2018). The orthologs of PL3 in *Aphelenchus avenae* and two *Bursaphelenchus* nematodes were grouped together with distinct clusters of *Meloidogyne* species (**fig. 5c**) is consistent with previous phylogenetic findings in PPNs (Danchin et al., 2010). The closest bacterial ortholog in the *Meloidogyne* clade was from *Curtobacterium flaccmfaciens* which is also known to cause bacterial wilt in the Fabaceae family (Júnior et al., 2012). Together, these results suggested some genes that were thought to play important roles in plant parasitism were in fact acquired earlier than the common ancestor of plant parasitic nematodes.

The majority of HGT gene families were of bacterial followed by fungal origin (**fig 4b**). We also identified genes that were acquired from non-bacterial donors in the last common ancestors of clade IV, as well as in more recent, different PPN lineages (**fig 4a**). This included the previously characterised fungal origin of GH45 (Kikuchi, Jones, Aikawa, Kosaka, & Ogura, 2004; Wu et al., 2016), This cellulase family is present in most Aphelenchoidea nematodes except *A. fujinensis* and *A. bicaudatus*. The GH16 family was independently acquired from a bacterial and fungal donor in the last common ancestor of clade IV nematodes and the *Aphelenchoides* genus, respectively (**fig 5b**).

Notably, we identified 40 orthogroups among PPNs that were transferred from the plant phylum Streptophyta, which is consistent with the finding of several sequences that are highly similar to plants in *H. glycine* (Elling et al., 2009) (**fig. 4b**). The closest plant orthologs included rice, maple and oak (**fig. 5d**) which are common hosts to many PPNs. Strikingly, of these orthogroups, 27 were present in *B. mucronatus* and enriched in the detoxification of cadmium and copper ion function **(table S9**), suggesting these genes may help *Bursaphelenchus* nematodes to degrade the toxin in pine wood hosts.

We identified 0.3-2.4%, 0.6-2.1% and 0.1-5.4% proteomes among Aphelenchoidea, Tylenchida and *Panagrolaimomorpha* nematodes that were predicted to be HGT (**fig. 4c**). The majority of these differences were the result of clade-specific evolution after speciation. The high copy number of HGT genes observed in *M. incognita* was a result of duplication (Szitenberg et al., 2017), indicated by the fact that the number of HGT orthologs of bacteria origin were over two times higher than any other species (**fig. S10**). The high number of HGT genes in *P. superbus* was consistent with a previous study (Schiffer et al., 2019) and likely to be species-specific.

To independently assess the accuracy of our approach and interrogate the fate of HGT genes, we constructed a phylogeny for every orthogroup containing identified HGT candidates. Members of Aphelenchoididae and Tylenchida orthologs in the majority of these orthogroups were predicted to be all HGT genes (with AI > 0; 54.6-76.5% vs. 77.3-89.4%). Genes from a species were typically grouped together in the orthogroup phylogeny regardless of being identified as HGT candidates, suggesting the genes that were not detected using our threshold shared common ancestries with those that were. Presumably, this was a result of accumulating substitutions over time.

Consistent with this observation, the more ancient acquired HGT orthogroups in PPNs contained higher copy numbers of these genes compared to recently acquired families (**fig.S11**). The instances included GH5 families with 12.5-70.6% of copies in Tylenchida that could not be identified as HGT candidates, suggesting duplication and possibly neo-functionalisation of the GH family in PPNs after being acquired from bacteria (**fig. 5a**). The differentiation was ongoing and observed in the *A. besseyi* species complex, which included the GH45 orthogroup with negative AI in two *A. oryzae* strains (**fig.S12**).

## Discussion

Characterising the diversity and comparing the genomes of plant parasitic nematodes has been of fundamental importance in understanding how such lifestyles arise and of practical importance in identifying candidate effectors and control methods. The latter has been addressed in several studies, focusing mainly on *Meloidogyne* (Grynberg et al., 2020). The *Aphelenchoides* genome assemblies presented in this study allowed us to gain a holistic view of the evolution of clade IV nematodes, which appeared to gain and lose many adaptations, including plant parasitism (Holterman et al., 2017). In their evolution, HGT genes have played important roles in functions related to these adaptations. The most recent comprehensive analyses of HGT in nematodes focused on plant parasitic nematodes (Grynberg et al., 2020) and found many of these genes were PPN-specific. The donors of these PPN-specific gene families were sympatric plant bacteria which may have facilitated the possibility of HGT (Danchin et al., 2010). Additional HGT events were identified in other clade IV nematodes (Danchin et al., 2016; Han et al., 2022; Schiffer et al., 2019; Wan et al., 2021; Zheng et al., 2016)

Our systematic investigation of HGT has shown that many of these families were acquired much earlier in the last common ancestor of clade IV. Sources of these donors may be symbionts like the case of insects (Xia et al., 2021), but currently nematode endosymbionts are restricted to *Wolbachia* and *Cardinium* (Brown et al., 2018) and were not identified in our analyses. Interestingly, many of the closest bacterial donors were from marine environments, raising the possibility that the last common ancestor of clade IV may have lived in a marine environment that underwent habitat transition (Holterman, Schratzberger, & Helder, 2019). However, we also identified donors of non-bacterial origin that were usually found in the environments that fit nematodes’ present day lifestyle. Now that more genome sequences are available, historical HGT events were detected in other nematode clades (Mayer et al., 2011; Zarlenga et al., 2019) as well as in the most recent common ancestor of major organism groups such as land plants (Ma et al., 2022), of moths and butterflies (Li et al., 2022), which contributed the hosts’ developmental roles and adaptations. These acquisitions were found to be episodic and likely took place in a time when either the host development or genome defence was vulnerable.

The successful delimitation of the *A. besseyi* species complex unambiguously into *A. oryzae* and the recently proposed *A. pseudobesseyi* has important implications in nematode management. Congruent delimitation was observed between genomes and 28S phylogenies, confirming the utility of species identification with existing molecular markers (Oliveira et al., 2019). We thus recommend that the taxonomic status of all *A. besseyi* strains be reclassified into either species. *A. besseyi* is generally believed to have limited mobility in natural habitats, so its lack of population structure in China (Xu et al., 2020) was suggested as a consequence of human-mediated dispersal. Our results also supported that *A. oryzae* appears to be more rice plant-specific compared to *A. pseudobesseyi* which was isolated more frequently in ornamental plants and other agronomic crops (Oliveira et al., 2019). A comprehensive collection across a wider geographical range and resequencing of strains previously designated as *A. besseyi* could confirm whether *A. oryzae* was responsible for all the white tip disease in rice plants and may lead to better characterisations of the biogeography and evolution of different cryptic species.

To conclude, the availability of the *Aphelenchoides* genome and our comparative analyses allowed us to pinpoint the major events of horizontal gene transfer in clade IV nematodes. The results have reinforced the importance of horizontal gene transfers contributing to multiple adaptations of these nematodes including plant parasitism. In addition, the various *A. besseyi* genomes will assist in developing molecular diagnostic tools to distinguish the specific diseases caused by the species complex.

## Methods

### DNA, RNA extraction and sequencing

Nematodes were cultured with *Alternaria citri* on PDA (potato dextrose agar) medium. All stages of nematodes were collected from the medium, washed with sterile distilled water, and purified by sucrose gradients. Genomic DNA was extracted using Qiagen Genomic-tip 100/G according to the manufacturer’s instructions, RNA extraction was conducted using Trizol, and then purified using a lithium chloride purification method. The DNA paired-end libraries were constructed using either a Nextera DNA Flex or KAPA hyper library prep kit (Illumina, San Diego, USA); the RNA paired-end libraries were constructed using a TruSeq Stranded mRNA library prep kit (Illumina, San Diego, USA). Both DNA and RNA pair-end followed with standard protocol and were sequenced by Illumina HiSeq 2500 (Illumina, USA) to produce 150-bp paired-end reads. The HiC library preparation was performed by Phase Genomics (Seattle, WA, USA) proximo HiC animal protocol with some modification in tissue processing. The enriched worms were finely chopped by microtube pellet pestle rods for about 2 minutes. The tissues were crosslinked by adding 1 ml crosslinking solution and incubate for 25 minutes with occasional mixing by rotation. 100 ul quenching solution was added to the crosslinked tissue and mixed for 20 minutes by rotation. The rest of the preparation steps follow the protocol. The library was sequenced by Illumina HiSeq 2500 (Illumina, USA) to produce 150-bp paired-end reads. APFT and AORT were using Pacbio sequencing system to produce long-read, and the rest of 4 *Aphelenchoides* strains (APVT, AORJ, *A. bicaudatus, A. fujianensis*) were sequenced using the Oxford Nanopore sequencing platform. The raw nanopore signals were basecalled by Guppy (Wick, Judd, & Holt, 2019) (ver 0.5.1) producing a total of 5.0-28.4 Gb sequences at least 1 kb in length.

### Assemblies of six *Aphelenchoides* spp

Raw reads of each species were assembled using Flye (ver 2.8.2)(Kolmogorov et al., 2019) assembler. The assemblies from Nanopore reads were corrected using Nanopore reads using Racon (Vaser, Sović, Nagarajan, & Šikić, 2017) (ver 1.4.6) and Medaka (ver 0.10.0; https://github.com/nanoporetech/medaka). All assemblies were further corrected using Illumina reads using Pilon (Walker et al., 2014) (ver 1.22) with five iterations. The *A. pesudobesseyi* VT assembly was scaffolded using HiC reads and subsequently curated in Juice-box (Durand et al., 2016) tools. The other five *Aphelenchoides* genomes were reference scaffolded based on this assembly using Ragtag (Alonge et al., 2019) (ver 1.1).

### Gene prediction and functional annotation

The identification of repetitive elements were computed by RepeatModeler (Flynn et al., 2020) (ver 1.0.8), TransposonPSI (ver 1.0.0; https://github.com/NBISweden/TransposonPSI) and USEARCH (Edgar & Bateman, 2010) (ver 8.1) based on the protocol by Berriman *et al*. (Coghlan, Coghlan, Tsai, & Berriman, 2018). Repeat locations were then identified by Repeatmasker (Tarailo-Graovac & Chen, 2009) (ver 4.0.9). RNA-seq reads of six *Aphelenchoides* strains were trimmed by Trimmomatic (Bolger, Lohse, & Usadel, 2014) (ver 0.36), and aligned to corresponding assemblies using STAR (Dobin & Gingeras, 2015) (ver 2.7.1a). From these mappings, transcripts were inferred using three approaches: i) assembled based on the mappings as guides using Trinity (Grabherr et al., 2011) (ver 2.84; option: default setting), reconstructed using ii) Stringtie (Pertea et al., 2015) (ver 1.3.4; option: default setting) and iii) CLASS2 (Song, Sabunciyan, & Florea, 2016) (ver 2.17; option: default setting). Transcripts generated from Trinity were realigned to the reference using GMAP (Wu & Watanabe, 2005) (ver 2017-11-15). The RNA-seq mappings were also used in BRAKER (Hoff, Lange, Lomsadze, Borodovsky, & Stanke, 2016) to train species parameter and generate an initial set of annotations. Proteomes of *Bursaphelenchus xylophilus* and *Caenorhabditis elegans* were downloaded from Wormbase ParaSite (WBPS14; Howe, Bolt, Shafie, Kersey, & Berriman, 2017) and used as homology guides to pick the best transcripts for each putative locus using MIKADO (Venturini, Caim, Kaithakottil, Mapleson, & Swarbreck, 2018) (ver 1.2.4; option: three Mikado steps, containing “prepare”, “serialize” and “pick” procedures), and were also used to train MAKER2. Finally, MAKER2 was invoked to generate a final set of gene annotations using picked EST evidence and protein evidence from MIKADO transcript and proteomes from closely related species (*Bursaphelenchus xylophilus* and *Caenorhabditis elegans*), and used gene models (BUSCO (Simão, Waterhouse, Ioannidis, Kriventseva, & Zdobnov, 2015), BRAKER, SNAP (Korf, 2004) and Augustus (Stanke et al., 2006)) as EST hints to train predicted data with three iterations. The APVT strain was predicted using trained models based on the manual curation of 975 genes.

### Comparative analyses

Proteomes of five plant-parasitic nematodes (*Bursaphelenchus xylophilus, Meloidogyne hapla, Meloidogyne incognita, Globodera pallida, Ditylenchus destructor*), two free-living nematodes (*Caenorhabditis elegans, Caenorhabditis briggsae*), six *Panagrolaimomorpha* (*Propanagrolaimus* sp. JU765, *Panagrellus revidius, Panagrolaimus superbus, Panagrolaimus* sp. PS1159, *Panagrolaimus davidi* and *Halicephalobus mephisto*) and one animal parasitic nematode (*Brugia malayi*) were downloaded from Wormbase WBP17 (Howe, Bolt, Shafie, Kersey, & Berriman, 2017) and the longest isoforms were selected for the further analyses. Only the longest isoform per gene was considered for subsequent analyses. Orthogroups were determined by Orthofinder (Emms & Kelly, 2019) (ver 2.2.7; options: -S diamond). Low-copy orthogroups were applied due to multiple genome duplication events in *M. incognita, A. avenae* and two *Panagrolaimus* species. Sequence alignments of each of the single-copy orthogroups were generated by MAFFT (ver 7.310; options: --maxterate 1000). Then, the concatenated alignment of all single-copy orthogroups was used to compute a maximum likelihood phylogeny using RAxML (Stamatakis, 2014) (ver 8.2.3; options: -s -T 32 -N 100 -f a -m PROTGAMMILGF) with 100 bootstrap replicates. Pfam copy numbers of all 27 nematodes were identified from the results of nematode proteomes blast against the database of Pfam website (ver 31; https://pfam.xfam.org/) using hmmscan (e value < 0.001; Finn, Clements, & Eddy, 2011). To identify putative effector enzymes, we searched the nematode proteomes against the CAZyme database (http://www.cazy.org; Drula et al., 2022) using hmmscan (Finn et al., 2011). We considered only sequences that were at least 80bp in length, had a conserved domain proportion of at least 0.35 of its length, and an e-value of less than 1e-15. Nigon elements of 15 rhabditid nematodes in *A. pseudobesseyi, B. xylophilus* and *C. elegans* were inferred based on (https://github.com/pgonzale60/vis_ALG; de la Rosa et al., 2021) which assigned Nigon units using BUSCO output (ver 4.8.4, options: genome; dataset: nematoda_odb10).

### Identification of the HGT genes

The probability of genes having been acquired via HGT was estimated by using Alienness Index (AI) (Rancurel et al., 2017). Our donor group were generated by non-Metazoans from NCBI nr database, and the recipient were Metazoans excluding the following species to prevent self-alignment: Aphelenchoidea, Tylenchida, Rhabditina, Spirurina and Cephaloboidea. The Alien Index (AI) was estimated by calculating the e-value of diamond (Buchfink, Xie, & Huson, 2014) (ver 2.0.14; option: blastx --evalue 0.001) best hits between the donor and recipient database. Orthogroups having at least one gene with an AI value over 30 were selected for further analysis. Gains and losses at each node were inferred using Phylip-Dollop (Campoy & González-Martín, 2017) (ver 3.69; options: fdollop -method d -ancseq). Some of the HGT family acquired branches were manually curated by their evolutionary place of gene phylogeny due to the fact that nematode genes with AI < 0 were clustered with other HGT genes. The highest AI value of nematode genes with classified taxonomy hit were chosen to represent the HGT origin in each orthogroup. Orthogroups with the same CAZyme annotated and nematode orthology gene AI higher than -50 in those Orthogroups were selected. AI < 0 genes were labelled with “*”. The orthologs were further combined with the HGT identified donor sequence from nr database and the specific cellulase sequence from CAZyme database. To reduce contamination, orthologs of Pfam domain were annotated and filtered by having at least one major domain (cellulase or pectate lyase).

Sequences of each HGT orthogroup were aligned using MAFFT (options: --maxiterate 1000 --genafpair) and trimmed by using trimAl (Capella-Gutiérrez, Silla-Martínez, & Gabaldón, 2009) (ver 1.4; options: -gappyout). The ortholog phylogenies were computed by using IQtree (Nguyen, Schmidt, Von Haeseler, & Minh, 2015) (ver 1.6.6; options: -bb 1000 -alrt 1000). For the CAZyme unclassified HGT orthogroups, the top 2 blast hits sequences from separated Uniprot (bacteria, fungi, land plants and insect) were used to confirm the HGT origin.

## AUTHORS CONTRIBUTION

IJT and PJC conceived the study. IJT led the study. YCL, TY and PJC sampled the *Aphelenchoides* nematodes. YiCL, HMK, WAL conducted the experiments. CKL analysed the data with input from HHL, IJT and CKL wrote the manuscript with input from YuCL, MRL, TY, TK and PJC.

## DATA AVAILABILITY

The sequencing data and annotation of six *Aphelenchoides* nematodes are publicly available in NCBI under the BioProject accession PRJNA834627. The accession numbers of individual assemblies are listed in **table S1** and scheduled in the WBPS18 of WormBase Parasite (https://parasite.wormbase.org/index.html). The accession numbers of individual samples are listed in **table S2**. The raw sequences, alignments and phylogenies of the HGT orthologs are available at https://github.com/lihowfun/CladeIV_HGT. The code used to perform this study is deposited at https://github.com/lihowfun/Aphelenchoides.git.

## Acknowledgement

We thank Pablo Gonzalez for his comments on the preprint version. We thank Christian Rödelsperger and the two reviewers for their comments. We would like to thank National Core Facility for Biopharmaceuticals (NCFB, 111-2740-B-492-001) and National Center for High-performance Computing (NCHC) of National Applied Research Laboratories (NARLabs) of Taiwan for providing computational resources and storage resources. This work was supported by Academia Sinica grant (AS-CDA-107-L01) to IJT.

