## Supplementary Info for "The *Aphelenchoides* genomes reveal substantial horizontal gene transfers in the last common ancestor of free-living and major plant parasitic nematodes"

### Supplementary Information Texts

#### Species choices

To investigate the genetic differences between members of the *A. besseyi* species complex, two *A. orazye* strains (RT and RJ) were collected from the rice field of Linnei, Nantou and Hiroshima, Japan; two *A. pseudobesseyi* strains (FT and VT) were isolated from bird's-nest fern in Mingjian, Nantou; and *Valliseria spiralis* was collected from Taichung aquarium. Two same genus (*A. bicaudatus* Fsh and *A. fujianesis* Dali) strains were isolated from ornamental nurseries of bird's-nest fern in Taiwan and a strawberry field in Dali, Taichung. To identify the *A. besseyi* species complex among samples, a PCR with different expected product size between *A. orazye* (926 bp) and *A. pseudobesseyi* (1,386 bp) was designed (5' TATGTCCGGAGTAAGTATTG and 3' TTAAACGAAAAGAATAAGCG) according to one-to-one ortholog in APFT and AORT.

#### 28S phylogeny

To confirm the relationship among members of the *A. besseyi* species complex, 28S sequences of *A. pseudobesseyi* (NCBI accession ID: KY510841.1, MH187564.1, KX356765.1, MT271870.1, MT271869.1 and MT271868.1)*, A. oryzae* (NCBI accession ID: KX356775.1, MK880169.1, KX356764.1, KY123694.1, KT692690.1, KX622689.1, KT692703.1, KX356776.1, KX356774.1, KY123697.1, MT271867.1 and KX356773.1), *Aphelenchoides gorganensis* (NCBI accession ID:KX357652.1) and *Aphelenchoides* ritzemabosi (NCBI accession ID: KX119133.1, KX356837.1 and KT692713.1) were collected from the reclassification of *A. besseyi* species complex study (Subbotin et al., 2020); then the download sequences were aligned to our six *Aphelenchoides* genomes using BLAT (Kent, 2002) (options: -t=dna -q=dna -out=blast9) to retrieve the 28S sequences among the assemblies. Then 28S phylogeny was produced by using RAxML with 100 bootstrap replicates.

#### Identification of telomere and telomeric genes

The telomere repeats in APVT and *B. xylophilus* genome were scanned by using Tandem Repeat Finder (ver 4.0.9). We then identified the telomere (TTAGGC) from the result. Neither this motif nor other motifs have been identified in the assembly of APVT, but the consensus motif has been found in nanopore raw reads. The alignment results showed it is likely to be trimmed due to the low read coverage in the assembler of Flye (ver 2.8.2) (Kolmogorov, Yuan, Lin, & Pevzner, 2019). However, the telomere motif TTAGGC was identified in four chromosomes of *B. xylophilus* assembly (Supplementary Table S3).

Telomeric protein-coding genes in nematodes were identified based on the homology genes in *C. elegans* (Fradin et al., 2017). The copy numbers were computed by comparison of proteins using diamond (Buchfink, Xie, & Huson, 2014) (ver 2.0.14; option: blastp –evalue 0.00001). The identity of homologs larger than 30 were selected and converted to a copy number matrix to plot a heatmap using pheatmap in R.

**Telomeric and meiosis genes**

We sought to identify possible causes of extensive fission and fusion in *A. peseudobesseyi*—the possibilities were any combination of telomere alteration, telomere maintenance and meiosis alteration. Orthologs of only 52 and one out of 98 telomere and ten meiosis-associated gene families in *C. elegans*, respectively, were identified in both the *Aphelenchoides* and the *Bursaphelenchus* genus (**Fig. S14**), suggesting either absence or divergence of these families may not be the primary reason for the karyotype rearrangement observed in *A. peseudobesseyi*. The majority of the ancestral sex chromosomes were unambiguously assigned to chromosome (chr) 2, and remapping of male sequences showed equal coverage along the chromosomes (**Fig. S15**), suggesting that the clade 10b nematodes exhibited a stochastic sex determination system that was recently characterized in *B. xylophilus* (Shinya et al., 2022).

### Supplementary Figures

**Fig. S1. The spatial proximity signal of HiC reads in *A. pseudobesseyi* VT (APVT) using Juicerbox** (Durand et al., 2016)**.** The self-blast against results is denoted with the red dots; the more red dots, the closer the genetic distance. The three blue blocks correspond to the three chromosomes in this strain.

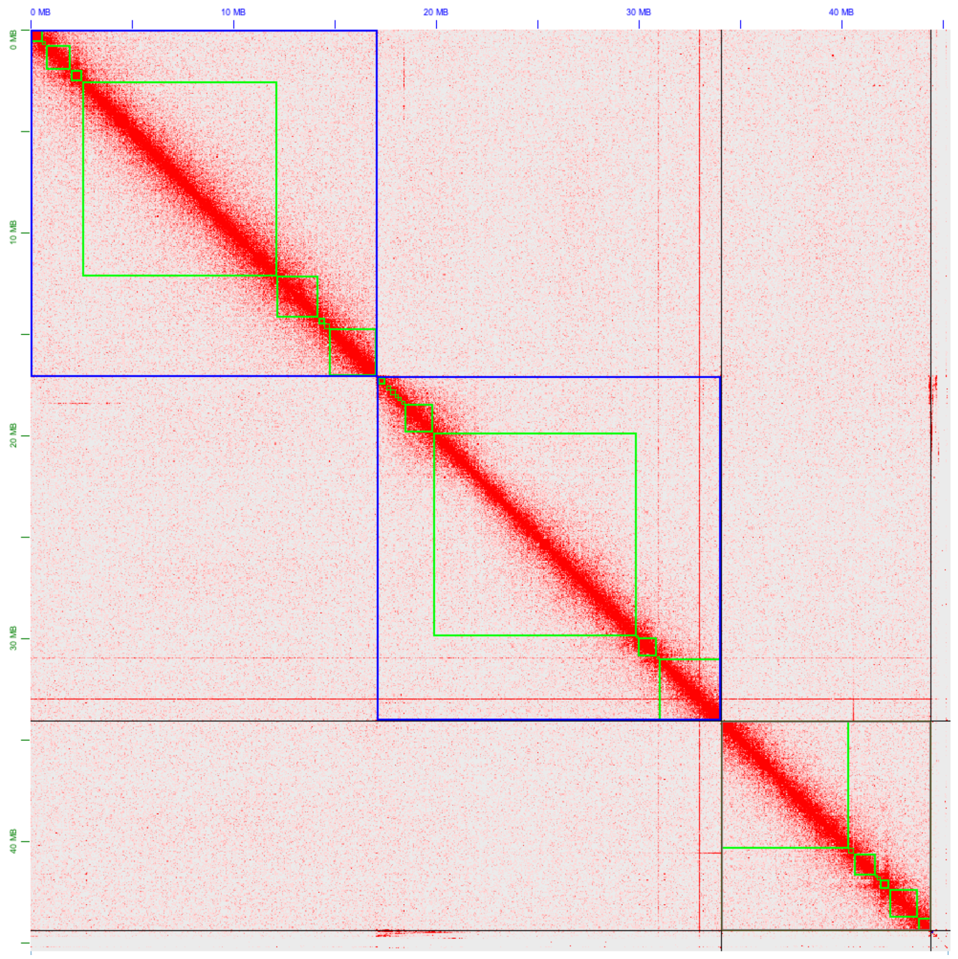

**Fig. S2. The polypoid prediction of *A. fujianensis* created using Smudge** (Ranallo-Benavidez, Jaron, & Schatz, 2019) Illumina paired end genomic sequences. The brighter colors indicate a higher number of kmer pairs. The Y-axis represents the total coverage of the kmer pairs, and the X-axis shows the normalized minor kmer coverage.

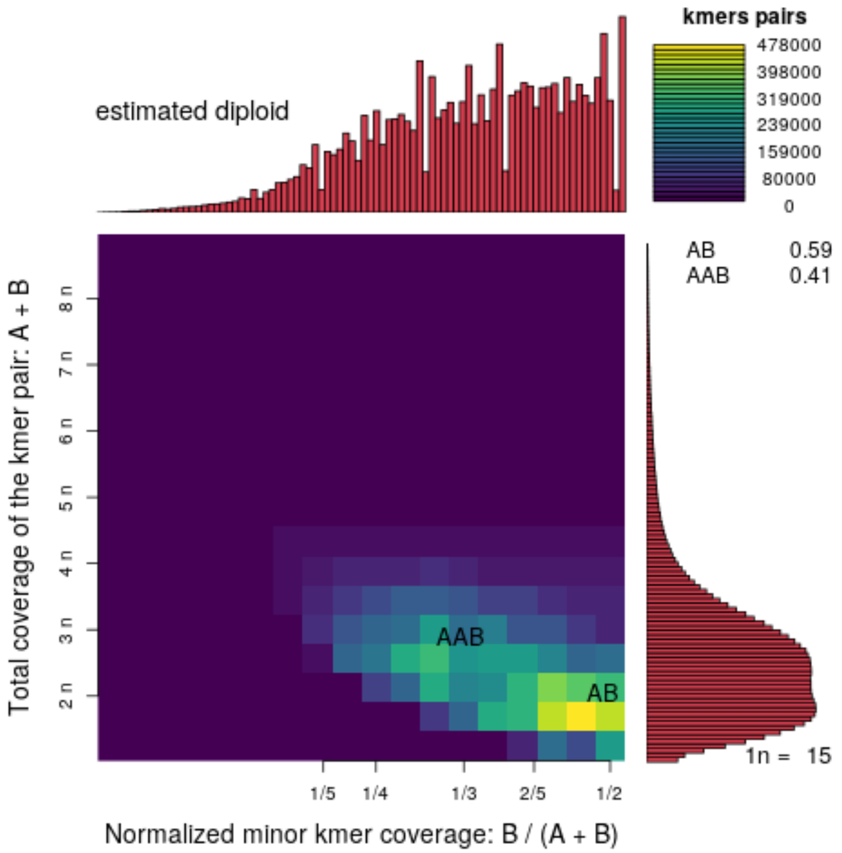

**Fig. S3. Separation of *Aphelenchoides* nematodes by using DA2 and DA3 of 28S.** Nodes with bootstrap support > 80% are labelled with grey circles. The *Aphelenchoides* species that were sequenced in this study are in bold.

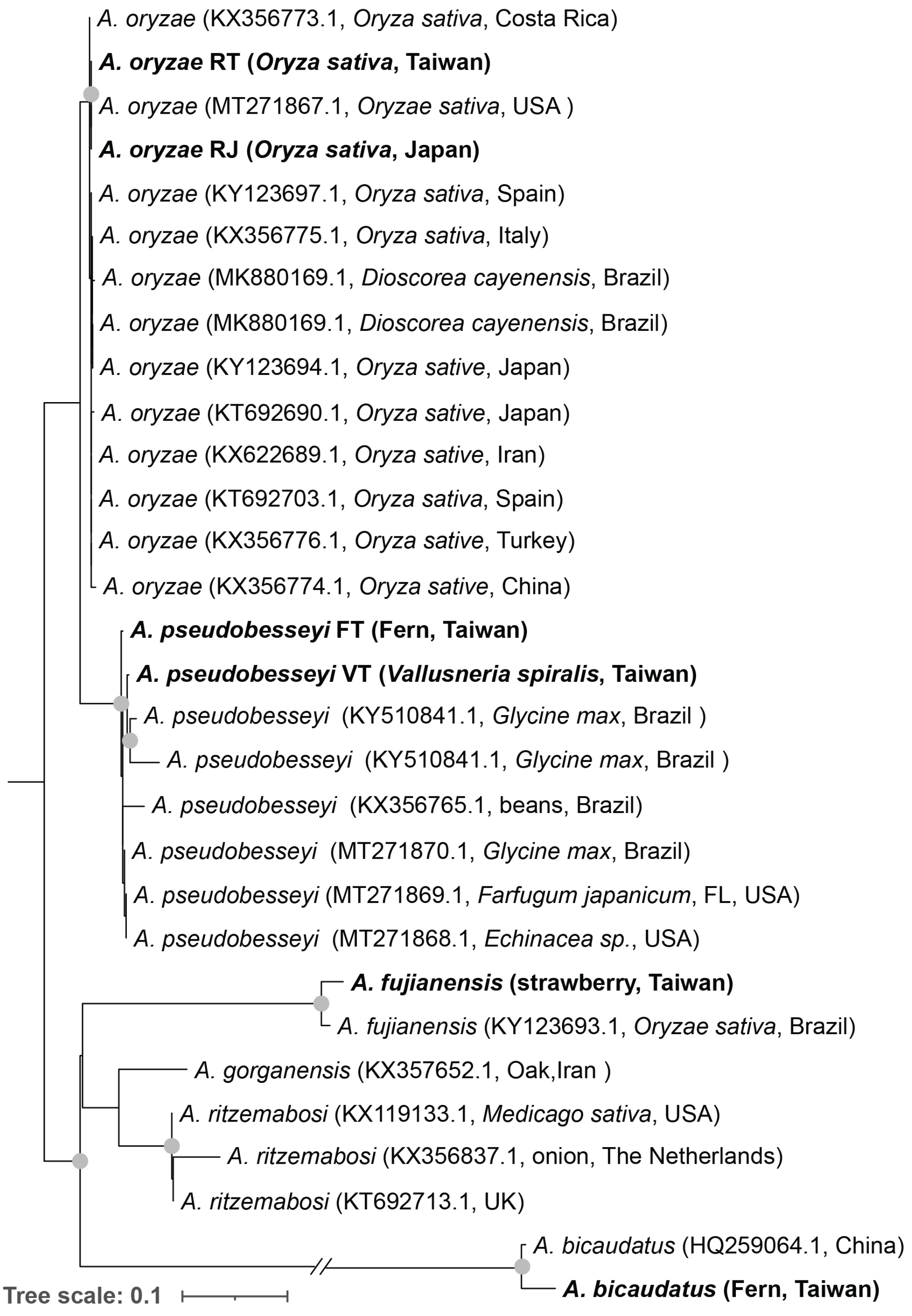

**Fig. S4. Whole genome median identity and protein similarity between members of *A. besseyi* species complex.** Pairwise protein similarity and nucleotide identity across four species in the *A. besseyi* complex are labelled and indicated by purple and red gradients, respectively.

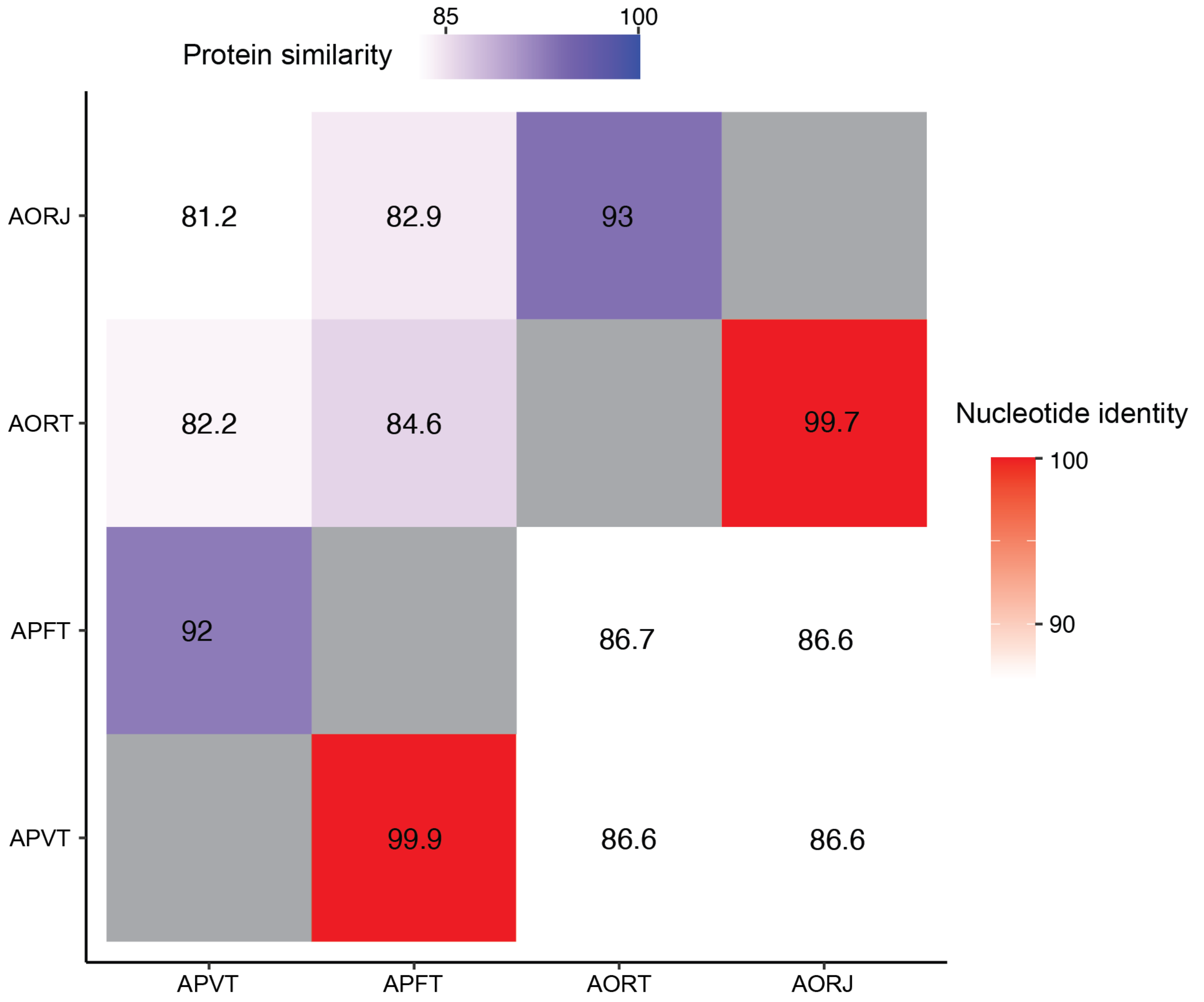

**Fig. S5. Prediction of past population size of *A. oryzae* RT (red) and *A. pseudobesseyi* VT (blue).**

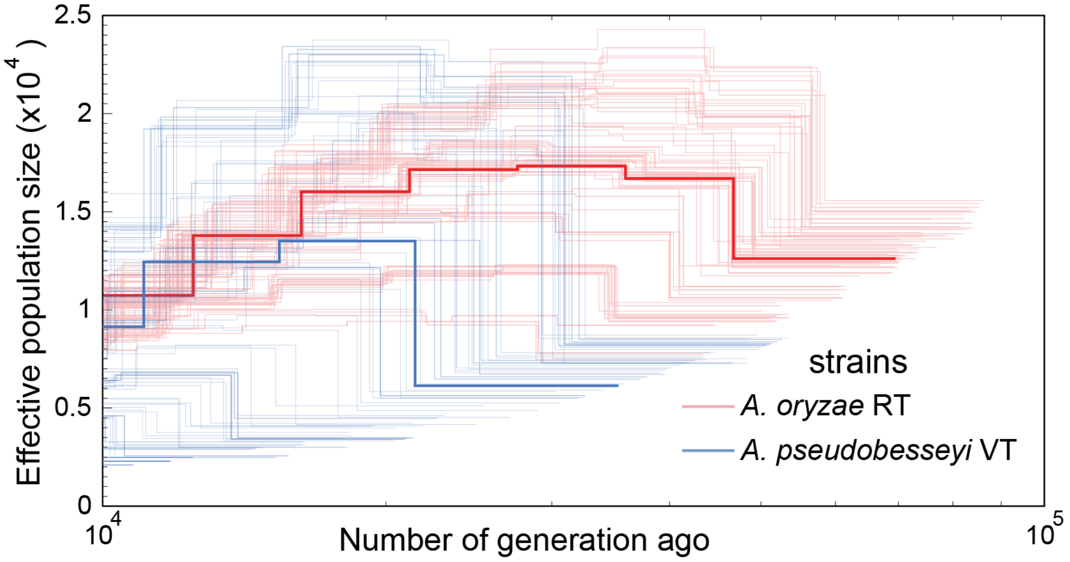


**Fig. S6. Distribution of transposable elements in the three chromosomes in *A. pseudobesseyi* VT and *A. oryzae* RT.** The DNA, LTR and LINE transposable contents in non-overlapping 10kb windows along the three chromosomes of strains are indicated by pink, green and yellow colours, respectively.

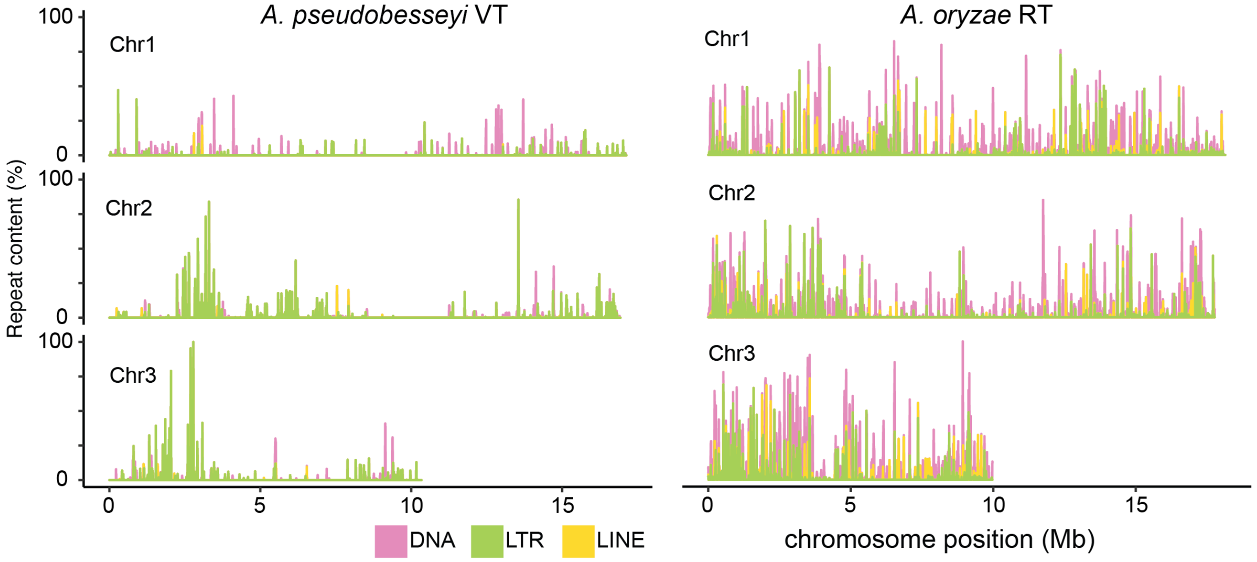

**Fig. S7. The coverage of *A. pseudobesseyi* VT male Illumina reads.** The reads depth of coverage of non-overlapping 10kb windows were normalized by the mean coverage of the longest chromosome.

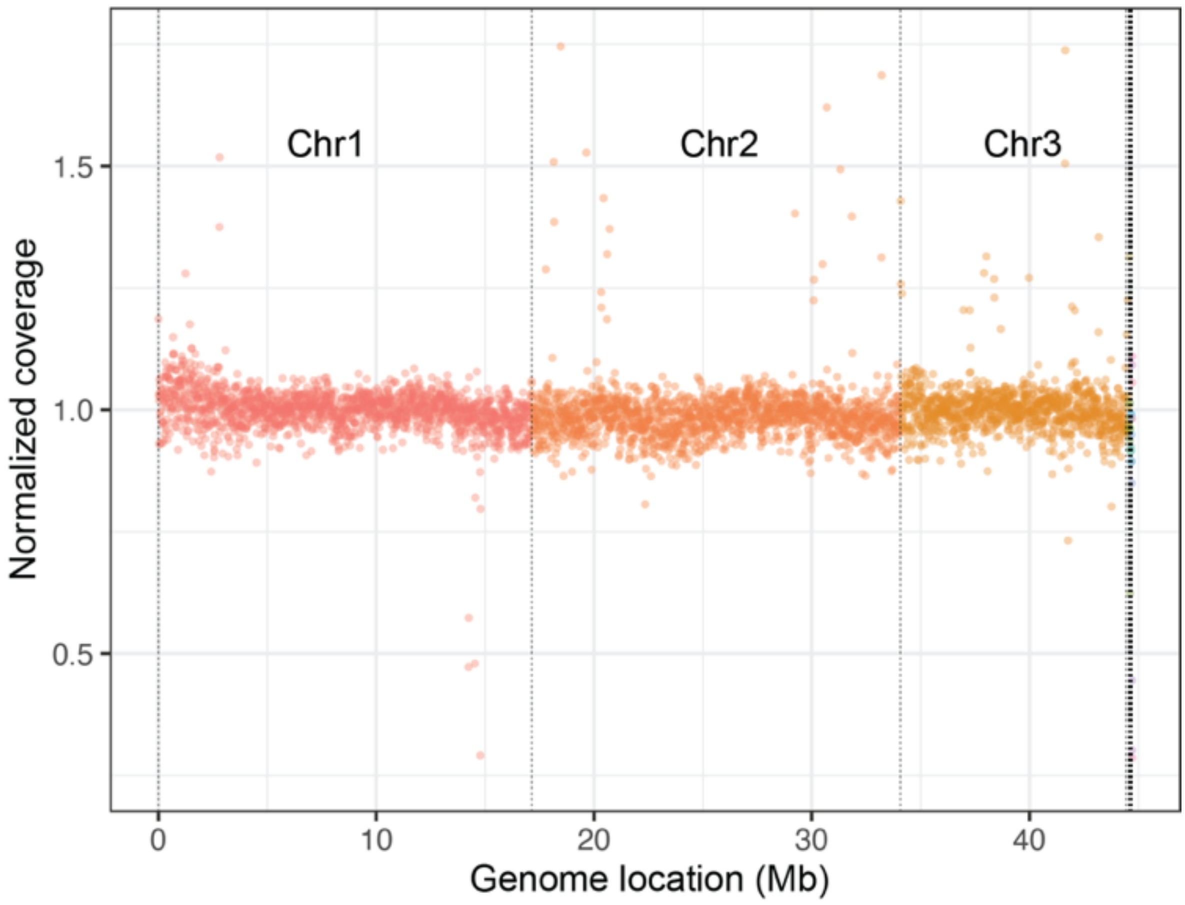

**Fig. S8. Phylogenetic relationship between GH32 copies in PPNs and *Panagrolaimomorpha* nematodes.** Different colours denote different kingdoms and species as shown in legend.

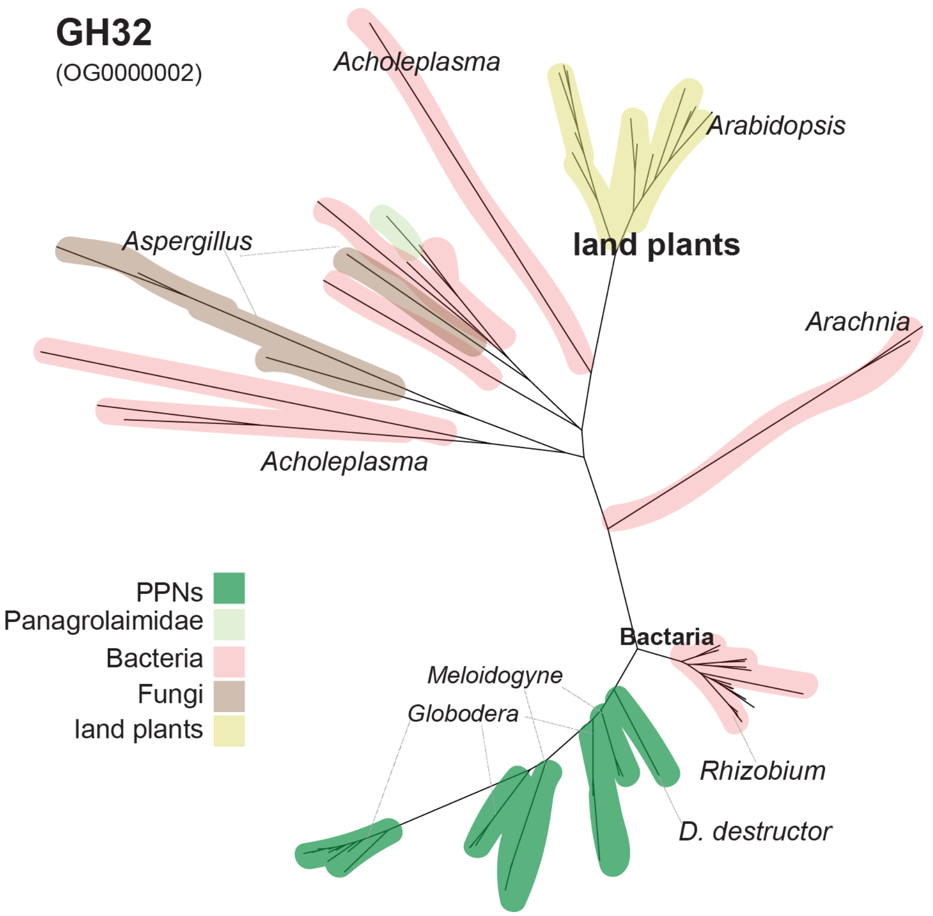

**Fig. S9. Phylogenetic relationship between GH43 copies in PPNs and *Panagrolaimomorpha* nematodes.** Different colours denote different kingdoms and species as shown in legend**.**

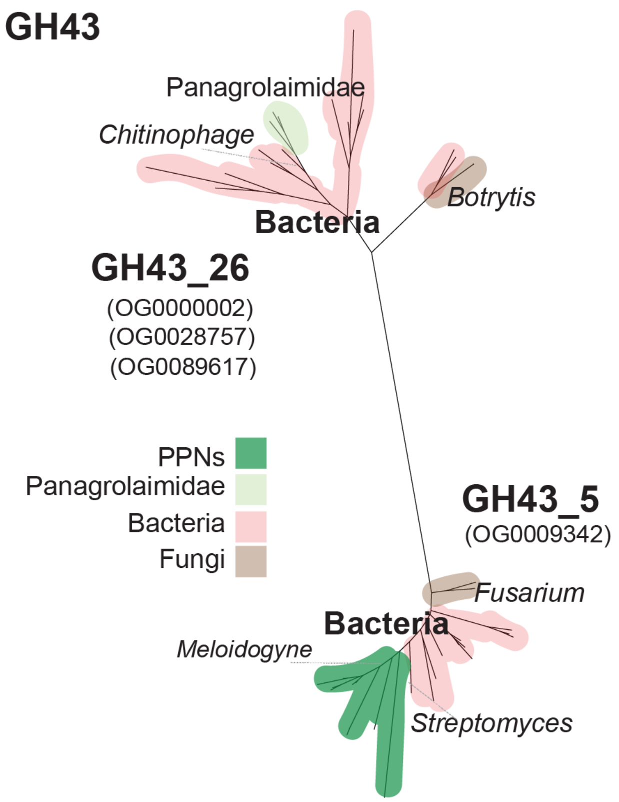

**Fig. S10. Total number of HGT events and genes across nematodes.** Different colours denote different phyla as shown in legend.

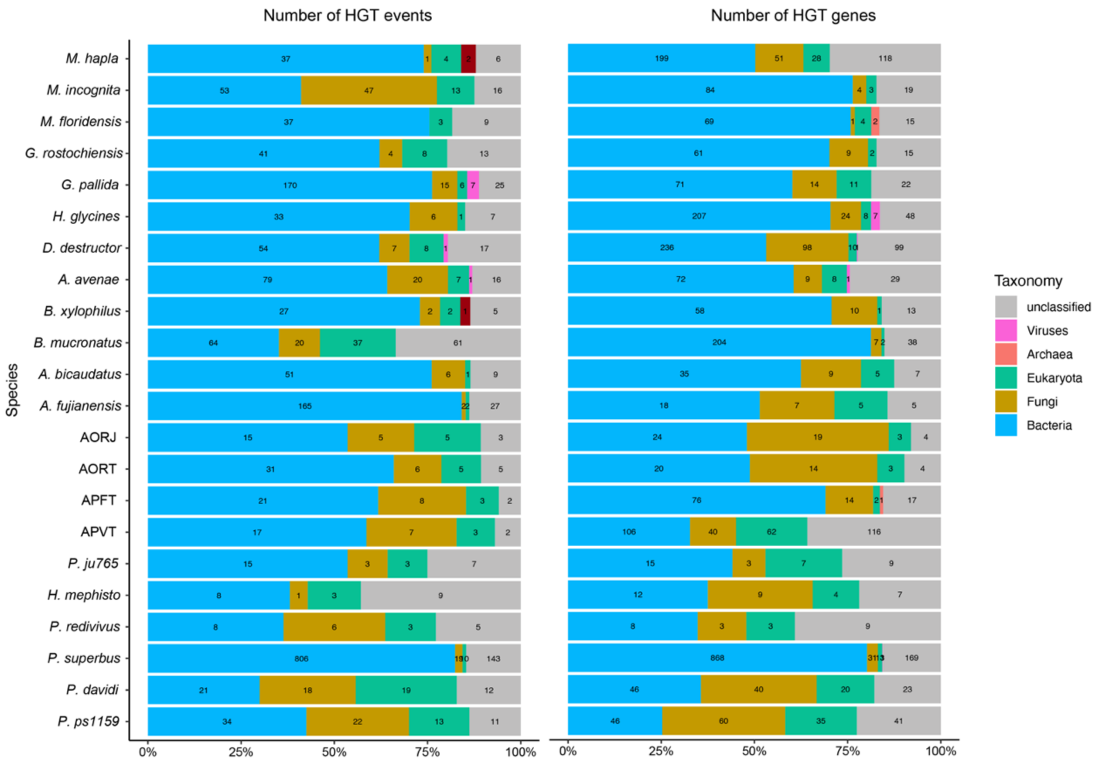

**Fig. S11. Proportions of genes in orthogroups identified as a HGT by Alienness in different nodes of the nematode phylogeny.** The HGT orthogroups containing 100%, 100-50% and 50-0% HGT copies are denoted with red, green and blue colours. The more ancient HGT families present a higher proportion of divergence copies.

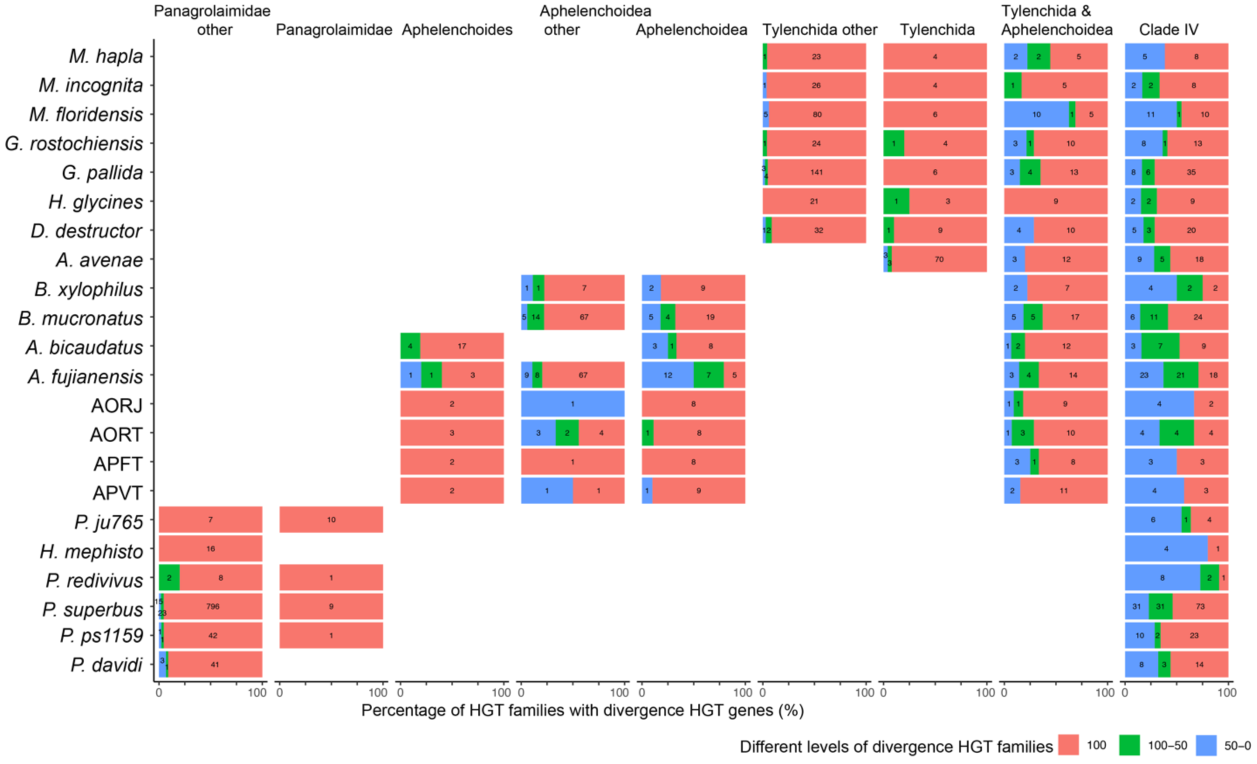

**Fig. S12. Phylogenetic relationship between GH45 copies in PPNs and their potential donor origin.** Different colours denote different kingdoms and species as shown in legend**.** Nematode gene copies with negative AI values are labelled with an asterisk. The HGT copies containing Pfam are labelled.

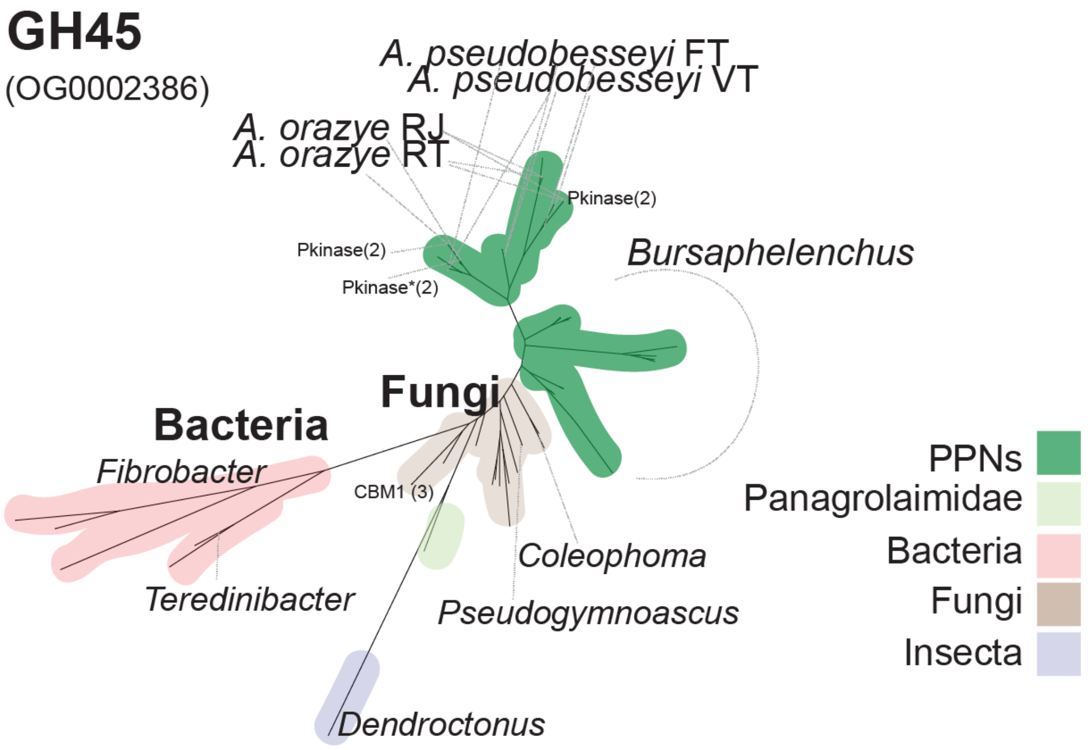


### Supplementary Tables

All tables are saved in a merged Excel xlsx file.
